# Skeletal muscle Nox4 knockout prevents and Nox2 knockout blunts loss of maximal diaphragm force in mice with heart failure with reduced ejection fraction

**DOI:** 10.1101/2022.05.27.493689

**Authors:** Ravi A. Kumar, Dongwoo Hahn, Rachel C. Kelley, Derek R. Muscato, Alex Shamoun, Nuria Curbelo-Bermudez, W. Greyson Butler, Svetlana Yegorova, Terence E. Ryan, Leonardo F. Ferreira

## Abstract

Patients with heart failure with reduced ejection fraction (HFrEF) experience diaphragm weakness that contributes to the primary disease symptoms of fatigue, dyspnea, and exercise intolerance. Weakness in the diaphragm is related to excessive production of reactive oxygen species (ROS), but the exact source of ROS remains unknown. NAD(P)H Oxidases (Nox), particularly the Nox2 and 4 isoforms, are important sources of ROS within skeletal muscle that contribute to optimal cell function. There are reports of increased Nox activity in the diaphragm of patients and animal models of HFrEF, implicating these complexes as possible sources of diaphragm dysfunction in HFrEF. To investigate the role of these proteins on diaphragm weakness in HFrEF, we generated inducible skeletal muscle specific knockouts of Nox2 or Nox4 using the Cre-Lox system and assessed diaphragm function in a mouse model of HFrEF induced by myocardial infarction. Diaphragm maximal specific force measured *in vitro* was depressed by ~20% with HFrEF. Knockout of Nox4 provided full protection against the loss of maximal force (*p* < 0.01), while the knockout of Nox2 provided partial protection (7% depression, *p* < 0.01). Mitochondrial respiration measured in permeabilized diaphragm muscle bundles increased with HFrEF or the knockout of Nox4 from skeletal muscle fibers (*p* < 0.05). Knockout of Nox2 from skeletal myofibers improved survival from 50 to 80% following myocardial infarction (*p* = 0.026). Our findings show an important role for skeletal muscle NAD(P)H Oxidases contributing to loss of diaphragm maximal force in HFrEF, along with systemic pathophysiological responses following myocardial infarction.

## Introduction

Heart failure with reduced ejection fraction (HFrEF) causes inspiratory dysfunction that contributes to the pathophysiology of the disease (Kelley & Ferreira, 2017). Patients with HFrEF display reduced inspiratory muscle strength and endurance that worsens with disease severity (Hammond *et al*., 1990; Mancini *et al*., 1995; Kelley & Ferreira, 2017; Miyagi *et al*., 2018). Maximal inspiratory pressure (MIP), a metric of inspiratory muscle strength, is an important indicator of clinical outcomes in patients with HFrEF: MIP is a better predictor of all-cause mortality than 6-minute walking distance and left ventricular ejection fraction (Ramalho *et al*., 2019), and patients that improved MIP following a cardiac rehabilitation program that included inspiratory muscle training had a lower risk of hospital re-admission or death than subjects that showed no improvement (Hamazaki *et al*., 2020). These studies highlight the importance of inspiratory muscle function and indicate that not all patients with HFrEF will respond to and benefit from inspiratory muscle training to improve inspiratory function (Hamazaki *et al*., 2020). Thus, further understanding of the mechanisms leading to inspiratory muscle dysfunction is critical for developing targeted treatments to improve mortality and quality of life in patients with HFrEF.

The diaphragm is the primary inspiratory muscle, and dysfunction in this skeletal muscle contributes to the inspiratory dysfunction seen in HFrEF. Muscle biopsies from patients and extensive work in animal models suggest contractile dysfunction underpins diaphragm abnormalities in HFrEF, along with reports of muscle atrophy, fiber-type shifts, and altered mitochondrial function (Howell *et al*., 1995; Tikunov *et al*., 1997; Ahn *et al*., 2015; Laitano *et al*., 2016; Adams *et al*., 2019; Coblentz *et al*., 2019; Kelley *et al*., 2020; Mangner *et al*., 2021). Reactive oxygen species (ROS) are key mediators of diaphragm dysfunction in HFrEF: increased markers of protein oxidation and intracellular redox shifts have been observed in the diaphragm of patients and animal models of HFrEF (Supinski & Callahan, 2005; Ahn *et al*., 2016; Laitano *et al*., 2016; Kelley *et al*., 2020; Mangner *et al*., 2021) but the source of excess ROS remains unknown.

NAD(P)H oxidases (Nox), specifically the Nox2 and 4 isoforms, are important sources of ROS in skeletal muscle (Sakellariou *et al*., 2013; Ferreira & Laitano, 2016; Henríquez-Olguín *et al*., 2019). Nox2 is activated by phosphorylation of its assembly subunit, p47^phox^, which coordinates the localization of several subunits to the sarcolemma and t-tubules (Ferreira & Laitano, 2016). Diaphragm biopsies from patients and animal models of HFrEF show increased expression of Nox2 subunits and phosphorylation of p47^phox^ (Ahn *et al*., 2015; Ahn *et al*., 2016; Mangner *et al*., 2021), implicating Nox2 as a primary source of ROS leading to diaphragm dysfunction. Wholebody knockout of p47^phox^ prevented the development of diaphragm contractile dysfunction in HFrEF (Ahn *et al*., 2015) but this protection could have originated from other cell types with high Nox2 activity such as immune cells (Winterbourn *et al*., 2016). Nox4, localized primarily in the mitochondria and sarcoplasmic reticulum (Ferreira & Laitano, 2016), is constitutively active and produces intracellular H_2_O_2_ dependent on its gene expression (Serrander *et al*., 2007). Nox4 protein content has recently been reported to increase in the diaphragm of patients with heart failure (Mangner *et al*., 2021). A recent report established a signaling axis in muscular dystrophy that involves crosstalk between Nox4 and Nox2 contributing to skeletal muscle weakness (Cully & Rodney, 2020). Increased Nox2 and 4 activity may also further ROS emission from mitochondria (Dikalov, 2011), which is evident in the diaphragm from rat models of HFrEF (Supinski & Callahan, 2005; Laitano *et al*., 2016; Coblentz *et al*., 2019), and cause post-translational modifications that impair mitochondrial respiration.

In the current study, we examined the role of skeletal muscle-specific Nox2 and Nox4 on diaphragm contractile properties, fiber size, and mitochondrial function (respiration and ROS) in a mouse model of HFrEF resulting from ischemic cardiomyopathy.

## Methods

### Animals

All procedures performed in animals followed guidelines established by the National Institutes of Health and were approved by the Institutional Animal Care and Use Committee of the University of Florida (Protocol #20189075). Animals were housed at our institution’s facilities under a 12-h:12-h light-dark cycle with access to standard chow and water *ad libitum*.

Inducible skeletal muscle-specific Nox2 knockout (skmNox2^KO^) male mice were generated by crossing mice carrying floxed Nox2 alleles (donated by Professor Ajay Shah (Sag *et al*., 2017)) with mice expressing an inducible Cre recombinase under the human skeletal actin (HSA) promoter (HSA-Cre) (McCarthy *et al*., 2012). The Cre recombinase is flanked by a mutated estrogen receptor (Mer) ligand-binding domain (MerCreMer) that permits inducible activation via tamoxifen injection. Inducible skeletal muscle-specific Nox4 knockout (skmNox4^KO^) male mice were generated by crossing mice carrying floxed Nox4 alleles (donated by Professor Junichi Sadoshima (Kuroda *et al*., 2010)) with mice expressing HSA-Cre. Once activated, the Cre recombinase identifies the loxP recognition sites flanking the Nox2 or Nox4 gene and excise that sequence in skeletal muscle. At approximately 8-10 weeks of age, mice received tamoxifen (5 consecutive days, 40 mg/kg body weight dissolved in ethanol, delivered in 100 μl sunflower oil) (skmNox2^KO^; skmNox4^KO^) or vehicle solution (100 μl sunflower oil) (skmNox2^+/y^; skmNox4^+/+^) through intraperitoneal injections (Saunders, 2011). Mice were randomly assigned to KO or control groups. The mouse diaphragm has a higher satellite cell content than limb muscle and sees a continuous contribution of satellite cells to the myonuclear pool over time, even under sedentary, non-stressed conditions (Keefe *et al*., 2015; Pawlikowski *et al*., 2015). To maintain knockout of Nox2 or Nox4 in the presence of ongoing satellite cell fusion over time, we additionally administered recombinant adeno-associated virus serotype 9 (rAAV9) containing HSA-Cre (to skmNox2^KO^ and skmNox4^KO^) or an empty vector (to skmNox2^+/y^ and skmNox4^+/+^) through a single intra-cardiac injection (~1 x 10^11^ vector genomes in 200 μl sterile saline) in the ventricular chamber. We anesthetized mice with an oxygen-isoflurane mixture (5% induction, 2-3% maintenance), removed hair from the abdomen and thorax, cleaned the area with chlorhexidine and sterile saline, and placed the mice in a supine position. A 31-guage needle containing the virus-saline mixture was inserted beneath the xyphoid process while applying gentle suction, angled cranially and slightly to the left of the sternum. Upon seeing blood return in the syringe, we slowly injected the virus.

Skeletal muscle-specific deletion of Nox2 was confirmed by polymerase chain reaction using primers from Sag et al. (2017). We extracted DNA from diaphragm, gastrocnemius, left ventricle, and liver tissue samples. Skeletal muscle Nox2 deletion was confirmed by amplification of a 225 base pair product that is present following recombination. Non-skeletal muscle tissues and skmNox2^+/y^ animals that did not receive tamoxifen displayed a 567 base pair product.

Skeletal muscle-specific deletion of Nox4 was confirmed by polymerase chain reaction based on the amplification of a ~1600 base pair product that forms after the excision of exon 9 of the Nox4 gene (forward primer, aacactgttggactcttcagacaca; reverse primer, ctcctgatgcatcggtaaagtc). Non-skeletal muscle tissues and skmNox4^+/+^ animals that did not receive tamoxifen do not display an amplified product, indicating that no recombination or gene excision had occurred.

Male mice were used in these experiments because of the higher incidence of HFrEF resulting from myocardial infarction in men (Virani *et al*., 2021). There is currently no scientific reason to expect differences between men and women regarding the development of diaphragm weakness as women diagnosed with HFrEF also experience inspiratory insufficiency, and female mice develop diaphragm weakness post-myocardial infarction (Adams *et al*., 2019).

### Surgery

Heart failure with reduced ejection fraction (HFrEF) was induced by myocardial infarction (MI) as previously described (Ahn *et al*., 2015; Coblentz *et al*., 2019). Four weeks after tamoxifen or vehicle injection and two weeks after viral administration, animals were anesthetized with a 5% isoflurane-oxygen mixture and placed on mechanical ventilation with 2-3% maintenance isoflurane. We then performed a left thoracotomy to expose the heart and ligated the left anterior descending coronary artery with 6-0 monofilament absorbable PGA suture (DemeSORB; Demetech). The ribs were approximated with 6-0 PGA suture, and the skin incision closed with 3-0 suture (DemeLON; Demetech). Sham surgeries were identical to the MI procedure but without ligation of the coronary artery. Mice were randomly assigned to sham or MI procedures within genetic strains.

Animals immediately received subcutaneous buprenorphine and topical bupivacaine. Buprenorphine injections were maintained every 8-12 hours for three days post-surgery. To minimize stress and enhance survival, animal cages were placed on a heating pad (set to maintain cage temperature at 32°C) and received supplemental medical-grade oxygen (0.35 F_i_O_2_) via a large animal O_2_ tent (Buster ICU; Kruuse; Langeskov, Denmark) for three days postprocedure.

### Echocardiography

Two to four weeks before terminal experiments, we assessed cardiac function by echocardiography as described previously (Ahn *et al*., 2015). Mice were maintained under ~2% isoflurane anesthesia (adjusted to keep heart rate between 400-500 bpm) while twodimensional M-mode ultrasound images were obtained in the parasternal short axis view (Aplio; Toshiba America Medical Systems, Tusin, CA). We determined left ventricular internal diameter during diastole (LVIDd) and systole (LVIDs) using Image J software. Fractional shortening (%) was calculated as (LVIDd – LVIDs) / LVIDd x 100. All echocardiography measures were completed between 17:00 and 21:00 h.

### Tissue harvesting and infarct size

Terminal experiments were conducted 14-16 weeks post-surgery and commenced between 08:30 h and 09:30 h. We anesthetized mice using isoflurane (5% induction, 2-3% maintenance) and performed a laparotomy and thoracotomy to collect tissue samples while the animals were in the surgical place of anesthesia.

We excised the diaphragm and heart and placed them in ice-cold bicarbonate-buffered solution (in mM: 137 NaCl, 5 KCl, 1 MgSO_4_, 1 NaH_2_PO_4_, 24 NaHCO_3_, and 2 CaCl_2_). Portions of the costal diaphragm were allocated for contractile, mitochondrial, and histological analyses. The remaining diaphragm was flash frozen in liquid nitrogen and stored at −80 °C for later biochemical assays.

Ventricles were separated, weighed, and the left ventricle was cut down the interventricular septum from the base to the apex to determine infarct size by planimetry (Finsen *et al*., 2005; Ahn *et al*., 2015). Inspiratory dysfunction worsens with the severity of HFrEF in patients (Kelley & Ferreira, 2017) and diaphragm dysfunction is prevalent in rodents with severe HFrEF (Kelley *et al*., 2020). For this reason, we restricted our HFrEF experimental groups to mice displaying severe HFrEF, defined as infarcted left ventricle area greater than 30% (Bayat *et al*., 2002) and right ventricle weight/body weight ratio two standard deviations greater than sham group average (indicative of right ventricular hypertrophy secondary to pulmonary arterial hypertension) (Lindsey *et al*., 2018; Kelley *et al*., 2020).

### Diaphragm contractile properties in vitro

We assessed diaphragm contractile properties in an isolated diaphragm strip as previously described (Kumar *et al*., 2020). A diaphragm strip was fixed between a glass rod and a dual-mode lever system (300C-LR; Aurora Scientific, Aurora, ON, Canada) and placed in a water-jacketed organ bath containing bicarbonate-buffered solution that was continuously gassed with 95% O_2_/ 5% CO_2_ at room temperature. Muscle contractile properties were assessed with a high-power biphasic stimulator (701c; Aurora Scientific) sending a supramaximal current (600 mA current; 0.25 ms pulse width; 0.3 s train duration) through two platinum electrodes flanking the muscle. We adjusted the length of the muscle to elicit maximal tetanic force (120 Hz) (optimal length; Lo) and increased the temperature of the bath to 37 °C. Following a ten-minute thermal equilibrium period, we measured diaphragm force at Lo during isometric contractions at frequencies of 1, 30, 50 and 300 Hz, separated by one minute rest intervals.

We measured muscle length at Lo using electronic calipers and determined muscle bundle weight. Force was normalized to muscle cross sectional area, calculated by muscle weight divided by the product of length and density of mammalian skeletal muscle (Close, 1972).

### Isolation and permeabilization of diaphragm fiber bundles for mitochondrial assessment

Diaphragm muscle bundles were prepared to assess mitochondrial function as recently described (Hahn *et al*., 2019). A portion of the right hemi-diaphragm was dissected with fine tweezers in ice-cold Buffer X (in mM: 7.23 K2EGTA, 2.77 Ca-K2EGTA, 20 imidazole, 20 taurine. 5.7 ATP, 14.3 PCr, 6.56 MgCl_2_-6H_2_O, 50 K-MES; pH 7.1) to remove branches of the phrenic nerve, and the abdominal layer of fascia and muscle fibers. The remaining pleural layer of fibers was then teased apart longitudinally to expose optimal surface area of fibers without inducing damage. We then permeabilized the bundles in Buffer X containing saponin (30 μg/ml) for 30 min at 4°C. Bundles were washed 3 × 5 min in buffer Z (in mM: 30 KCl, 10 KH_2_PO_4_, 5 MgCl_2_-6H_2_O, 105 K-MES, and 0.5 mg/ml BSA, pH 7.1), after which we proceeded with assessment of mitochondrial respiration or substrate-induced H_2_O_2_ emission.

### Mitochondrial respiration

We performed high-resolution O_2_ respirometry at 37°C in buffer Z with 20 mM creatine monohydrate and 10 μM Blebbistatin using an O_2_K Oxygraph (Oroboros, Innsbruck, Austria). We used a substrate-inhibitor titration protocol to measure oxygen flux (*J*O_2_) under the following incremental conditions: 0.5 mM malate + 10 mM glutamate, ADP (0.5 mM, 8 mM), 10 mM Cytochrome C, 10 mM succinate, and 10 μM rotenone. Oxygen concentration in the assay buffer was brought up to ~ 400 μM at the start of each experiment and re-oxygenated when O_2_ content dropped below 250 μM. Bundles were immediately blotted dry, flash frozen in liquid nitrogen, and stored at −80°C for later measurements of total protein and immunoblotting experiments. Oxygen flux (*J*O_2_, pmol/s) was normalized by bundle total protein. Two bundles from the same animal were run simultaneously and the results were averaged for statistical analysis. Respiratory control ratio (RCR) was calculated as the ratio of state III (maximal ADP stimulated; 8mM ADP) to state II respiration (malate + glutamate). Bundles were excluded from final analyses if there was a >10% increase in *J*O_2_ following the addition of cytochrome C, indicating fiber bundle damage.

### Mitochondrial H_2_O_2_ emission

Substrate induced H_2_O_2_ emission was assessed in permeabilized diaphragm fiber bundles using a fluorometer (λ_excitation_ = 565 nm, λ_emission_ = 600 nm, Fluorolog-3; Horiba Jobin Yvon, Edison, NJ) as previously described (Coblentz *et al*., 2019; Hahn *et al*., 2019). Bundles were placed into a quartz cuvette containing Buffer Z with 10 μM Amplex Ultra Red (Life Technologies, Eugene, OR), 25 μM Blebbistatin, 1mM EGTA, and 1 U/ml horseradish peroxidase that was heated to 37 °C and continuously stirred with a magnetic stir bar. We measured Amplex Ultra Red fluorescence under baseline conditions followed by the addition of 10 mM succinate. Amplex Ultra Red fluorescence was converted to H_2_O_2_ emission (*J* H_2_O_2_) using a standard curve that was determined after each experiment. Bundles were immediately blotted dry and weighed for data normalization (*J* H_2_O_2_: pmol/s/mg WW).

### rtPCR

We isolated RNA from tissue samples using the Direct-Zol RNA Microprep kit from Zymo Research (Irvine, CA). Samples were homogenized in TRI-Reagent (T9424; Sigma Aldrich) using stainless steel beads and a bullet blender (BBY24M; Next Advanced, Troy, NY). We assessed RNA quantity and quality via UV spectroscopy (Nanodrop 2000; ThermoFisher), then generated cDNA using the Superscript IV First-Strand synthesis system (18091050; Invitrogen). Real time PCR was performed on a Quantstudio 3 thermocycler (Thermo Fisher) using Taqman Universal Master Mix II (4440040; ThermoFisher) and Taqman probes (all from Thermo Fisher) for Nox4 (Mm00479246_m1), Nox2 (*CYBB*; Mm01287743_m1) and p47p^hox^ (*NCF1*; Mm00447921_m1). Results using Taqman probes were normalized to *HPRT* (Mm00446968_m1). Because the Taqman probes used in these experiments target exons that are still transcribed following genetic recombination, we completed additional experiments using SYBR Green (4309155; ThermoFisher) and primers specifically targeted to areas of recombination in the Nox2 and Nox4 models. For detecting Nox2 in the Nox2 floxed model, we used primers targeted to exons 1 and 2 (F: AGA GAG GCA GAA CCA ACA CT; R: CCC CAA CCA CAC CAG AAT GA). For detecting Nox4 in the Nox4 floxed model, we used primers targeted to exons 8 and 9 (F: GCT CAT TTC CCA CAG ACC TGG; R: GGT GAC AGG TTT GTT GCT CCT). These results were normalized to HPRT expression (F: CTC ATG GAC TGA TTA TGG ACA GGA C; R: GCA GGT CAG CAA AGA ACT TAT AGC C). Gene expression was calculated relative to the skmNox4-sham or skmNox2-sham group using the ΔΔC_T_ method.

### Gel electrophoresis and immunoblotting

To assess protein abundance via immunoblotting, diaphragm samples were homogenized 1:10 w:v in RIPA Buffer (50 mM Tris [pH 7.5], 150 mM NaCl, 0.075% NaDoC, 1% IGEPAL CA-60) with 1x Halt protease and phosphatase inhibitors (78440; ThermoFisher) on ice using Dual Kontes glass tissue grinders. We centrifuged the homogenates (16,000 x g, 15 min, 4 °C), then determined the protein content of the resulting supernatant using the DC Assay (Bio-Rad). Supernatants were then diluted 3:1 with 4x Laemmli sample buffer (125 mM Tris-HCl [pH 6.8], 20% glycerol, 4% SDS, 10% 2-mercaptoethanol, and 0.02% bromophenol blue).

We prepared additional tissue homogenates using diaphragm fiber bundles from mitochondrial respiration experiments to assess differences in mitochondrial electron transport system complexes and citrate synthase content. We homogenized fiber bundles in 40 μl of 1x Laemli Buffer (Bio-Rad) containing 0.35 M DTT at room temperature using a Dual Kontes plastic tissue homogenizer attached to an electric drill (30 s, high torque setting). Homogenates were centrifuged at 1,000 x g for 10 min at room temperature and the resulting supernatant was transferred to a fresh tube. We determined protein content of the lysate via gel electrophoresis (described below) using a 3-point bovine serum albumin (BSA) standard curve. Total protein content of each individual bundle was determined by multiplying the calculated protein content (μg/μl) by 40 μl, the initial dilution volume.

For gel electrophoresis, we ran similar amounts of protein per lane (~10-20 μg, depending on target) on a 4-20% Criterion TGX stain-free gel (Bio-Rad) at 200 V for 50 min with the cassette surrounded by ice. Gels were activated and scanned using a Gel Doc EZ imager (Bio-rad) for determination of total protein per lane, after which proteins were transferred onto a nitrocellulose membrane at a current of 200 mA for 2 hours at 4 °C. Following transfer, the membrane was washed for 5 minutes in TBS, then blocked in Intercept blocking buffer (LI-COR, Lincoln, NE) for one hour at room temperature. We then incubated membranes in primary antibodies diluted in blocking buffer with 1% TWEEN-20 overnight at 4 °C. Primary antibodies included citrate synthase (1:1,000 dilution ab96600; Abcam, Cambridge, MA), OXPHOS antibody cocktail (1:1,000 dilution ab110413; Abcam), and RyR1 (1:500 dilution MA3-925; Invitrogen). Membranes were washed 4×5 min in TBS-T, then incubated with appropriately conjugated secondary antibodies (IRDye_800_ 1:20,000 dilution; IRDye_680_ 1:40,000 dilution – LI-COR) for 1 hour at room temperature. After 4×5 min additional washes in TBS-T and 1×5 min rinse in TBS, membranes were scanned using an Odyssey Infrared Imaging System (LI-COR).

We used Image Lab 5.0 software (Bio-Rad) to quantify the total protein signal in each gel lane after gel electrophoresis. We quantified the immunoblot signal of target proteins using Image Studio Lite (LI-COR, Lincoln, NE). The immunoblot signal of each target protein was then normalized by the total protein signal measured in the corresponding lanes.

We tested the linear range for all antibodies used in this study using pooled samples and loaded protein contents that were approximately in the mid-portion of the linear range of detection for each antibody. Each gel and membrane included an abbreviated 3-point linearity to further confirm that individual samples were loaded within the linear range of detection.

### Fiber type distribution and cross-sectional area

A diaphragm bundle was embedded in Tissue-Tek OCT freezing medium, frozen in liquid-nitrogen cooled-isopentane, and stored at −80 °C. Later, we prepared 10 μm cross-sections of diaphragm bundles using a cryostat (Leica, CM 2050S model) cooled to approximately −20°C, transferred sections to frosted microscope slides, and followed a previously described protocol to stain for myosin heavy chain isoforms (Kelley *et al*., 2020). Briefly, we incubated slides in 1:2000 wheat germ agglutinin (WGA) Texas Red (Molecular Probes) and primary antibodies specific for myosin heavy chain Type I (A4.840, 1:15; Developmental Studies Hybridoma Bank) and Type IIa (SC-71, 1:50; Developmental Studies Hybridoma Bank). Sections were then incubated with fluorescently conjugated secondary antibodies (Goat x Mouse IgM Alexa 350 and Goat x Mouse IgG Alexa 488, Invitrogen) and imaged using an inverted fluorescence microscope (Axio Observer, 10x objective lens) connected to a monochrome camera (Axio MRm) and controlled with Zen Pro software (Carl Zeiss Microscopy). We used the high throughput semi-automatic Myovision software (Wen *et al*., 2018) to quantify fiber cross sectional area and detect fiber types based on antibody detection. Fibers that were not detected by Type I or IIa antibodies were labeled as Type IIx.

### Glutathione (GSH) content and redox state

We determined glutathione content and redox state using high-performance liquid chromatography as described previously (Jones, 2002; Ferreira *et al*., 2009). Diaphragm samples were homogenized in perchloric acid with 0.2 M boric acid and 10μM γ-glutamylglutamate using Dual Kontes tissue grinders, then sonicated and stored at −20 °C overnight. We then centrifuged the homogenates (16,000 g, 20 min, 4 °C) and stored the supernatant at −80 °C. After determining the protein content of the resulting pellet using the DC Assay (BioRad), we sent frozen supernatants for determination of GSH, CysGSH, and GSSG at the Emory Clinical Biomarkers Laboratory (Emory University, Atlanta, GA). Results were normalized by total protein content of the pellet. Oxidized glutathione content was determined by the sum of CysGSH content and 2 x GSSG content.

### Statistics

Data in text and tables are presented as mean ± SD. Individual data (scatterplots) and group means (bars) are shown in figures unless stated otherwise. We performed tests for normality (Shapiro-Wilk) and equal variance (Brown-Forsythe), and log-transformed non-parametric data before running parametric statistical tests. We analyzed effects and interactions of surgical treatment (*factor 1*: sham vs. HFrEF) and genetic strain (*factor 2*: skmNox2/4^+/+^ vs. skmNox2/4^KO^) by two-way ANOVA. When there was a statistically significant interaction, we followed up with Bonferroni’s test for post hoc comparisons (SigmaPlot v. 14.0; Systat Software, San Jose, CA). In select cases, we performed post-hoc analyses when the interaction effect did not reach significance based on previous recommendations (Wei *et al*., 2012). We used unpaired Student’s *t*-test (parametric) to compare infarct size between strains. For assessing differences in fiber cross-sectional area, we applied a linear mixed model analysis (SPSS v26 IBM, Armonk, NY) because of the variable number of fibers measured for each animal. Survival rates post-MI were compared using the Gehan-Breslow-Wilcoxon test. This test places greater weight on differences in survival at earlier time points if hazards ratios are not consistent over time, as is the case with mortality post-MI (Lindsey *et al*., 2018). All statistical tests were conducted using two-tailed tests and statistical significance declared when *p* < 0.05. Wherever feasible, we report exact *p* values and have taken recent recommendations into consideration for our data interpretation (Wei *et al*., 2012; Amrhein *et al*., 2019; Curran-Everett, 2020).

## Results

### Animals

All animals included in final analyses displayed the anticipated PCR amplified products, indicating appropriate genetic strain and presence (KO) or absence (+/+ for Nox4, +/y for Nox2) of recombination by Cre recombinase (Fig. S1 A, B). Gene expression of *Nox2* and *NCF1* decreased with skeletal muscle knockout of Nox4, regardless of surgical treatment (Fig. S1 D, E), whereas *Nox4* expression increased with HFrEF but showed no change with Nox4^KO^ compared to vehicle injected shams (Fig. S1 C). In skmNox2^+/y^ groups, Nox2 content decreased by 50% following genetic recombination, while HFrEF caused a decrease in *Nox4* and *Nox2* gene expression (Fig. S1).

Characteristics of experimental animals are listed in Tables 1 and 2. For investigations into the effect of Nox4 on diaphragm function in HFrEF, a total of 76 animals underwent the MI procedure. Post-operative survival rate for skmNox4^+/+^ and skmNox4^KO^ were not different (Fig. 1A). After applying infarct size and right ventricle/body weight ratio inclusion criteria for severe HFrEF, seven skmNox4^+/+^ (five excluded) and six skmNox4^KO^ (five excluded) HFrEF animals were used in final analyses of diaphragm function. The skmNox4^+/+^ and skmNox4^KO^ sham surgery groups each contained seven animals. Thus, the total number of animals used for diaphragm analysis was n = 27

**Figure 1.**
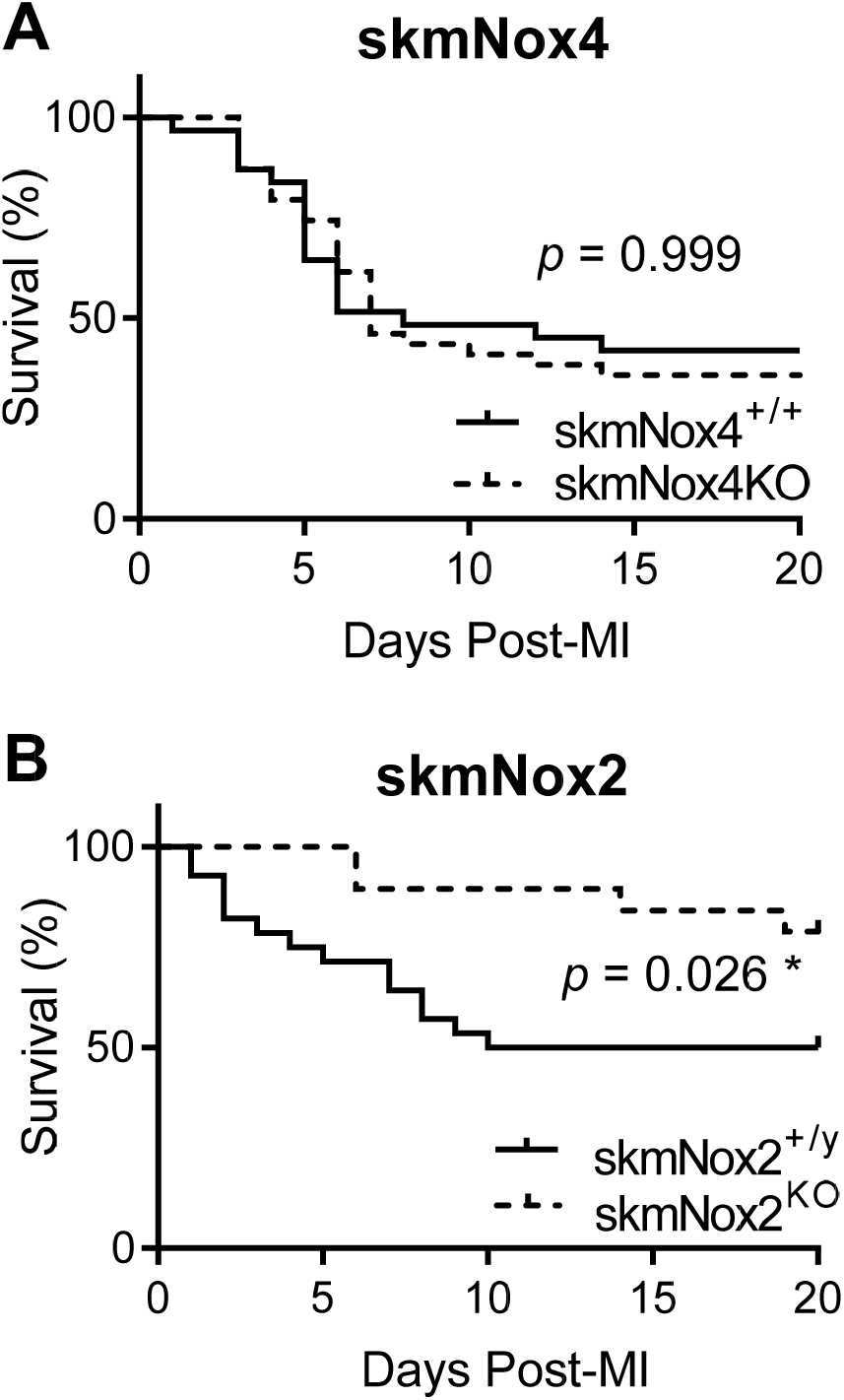
Kaplan Meyer survival curves post-myocardial infarction. Knockout of skeletal muscle Nox2 (B) but not Nox4 (A) improves survival post-myocardial infarction (MI). *P* values are from Gehan-Breslow-Wilcoxon test. \**p* < 0.05.

**Table 1.** skmNox4 - Animal characteristics.

|  | SkmNox4<sup>+/+</sup> |  | SkmNox4<sup>KO</sup> |  | *p* values |  |  |
| --- | --- | --- | --- | --- | --- | --- | --- |
|  | Sham (n = 7) | HFrEF (n = 7) | Sham (n = 7) | HFrEF (n = 6) | Surgery | Nox4 | Interaction |
| Terminal body weight (g) | 31.46 ± 2.7 | 30.9 ± 1.7 | 29.9 ± 3.1 | 30.5 ± 2.1 | 0.995 | 0.315 | 0.581 |
| Fractional Shortening (%) | 30.0 ± 4.6 | 13.5 ± 2.6 | 25.4 ± 5.5 | 15.5 ± 4.3 | < 0.001 * | 0.473 | 0.067 |
| LVIDD (cm) | 0.45 ± 0.04 | 0.75 ± 0.10 | 0.48 ± 0.05 | 0.67 ± 0.13 | < 0.001 * | 0.423 | 0.085 |
| LVIDS (cm) | 0.32 ± 0.04 | 0.65 ± 0.08 | 0.36 ± 0.05 | 0.57 ± 0.14 | < 0.001 * | 0.587 | 0.072 |
| Heart Rate (BPM) | 424 ± 27 | 465 ± 66 | 443 ± 44 | 426 ± 67 | 0.567 | 0.637 | 0.176 |
| Infarct size (%) | - | 39 ± 5.7 | - | 42 ± 8.9 |  | 0.529 |  |
| RV weight (mg) | 28 ± 2.8 | 48 ± 11.0 | 27 ± 2.8 | 44 ± 8.7 | < 0.001 * | 0.331 | 0.555 |
| LV weight (mg) | 105 ± 8.7 | 188 ± 35.6 | 108 ± 12.6 | 202 ± 71.4 | < 0.001 * | 0.665 | 0.953 |
| RV weight/body weight | 0.88 ± 0.04 | 1.57 ± 0.38 | 0.89 ± 0.11 | 1.44 ± 0.26 | < 0.001 * | 0.611 | 0.506 |
| LV weight/body weight | 3.33 ± 0.15 | 6.10 ± 1.21 | 3.63 ± 0.43 | 6.7 ± 2.7 | < 0.001 * | 0.414 | 0.871 |

Data are presented as mean ± SD. Statistical analysis by Two-way ANOVA with Bonferroni's post-hoc test when appropriate. Infarct size compared using Student's *t*-test. HFrEF, heart failure with reduced ejection fraction; LVID, left ventricular internal diameter during diastole (D) or systole (S); RV, right ventricle; LV, left ventricle. \* *p* < 0.05.

**Table 2.**
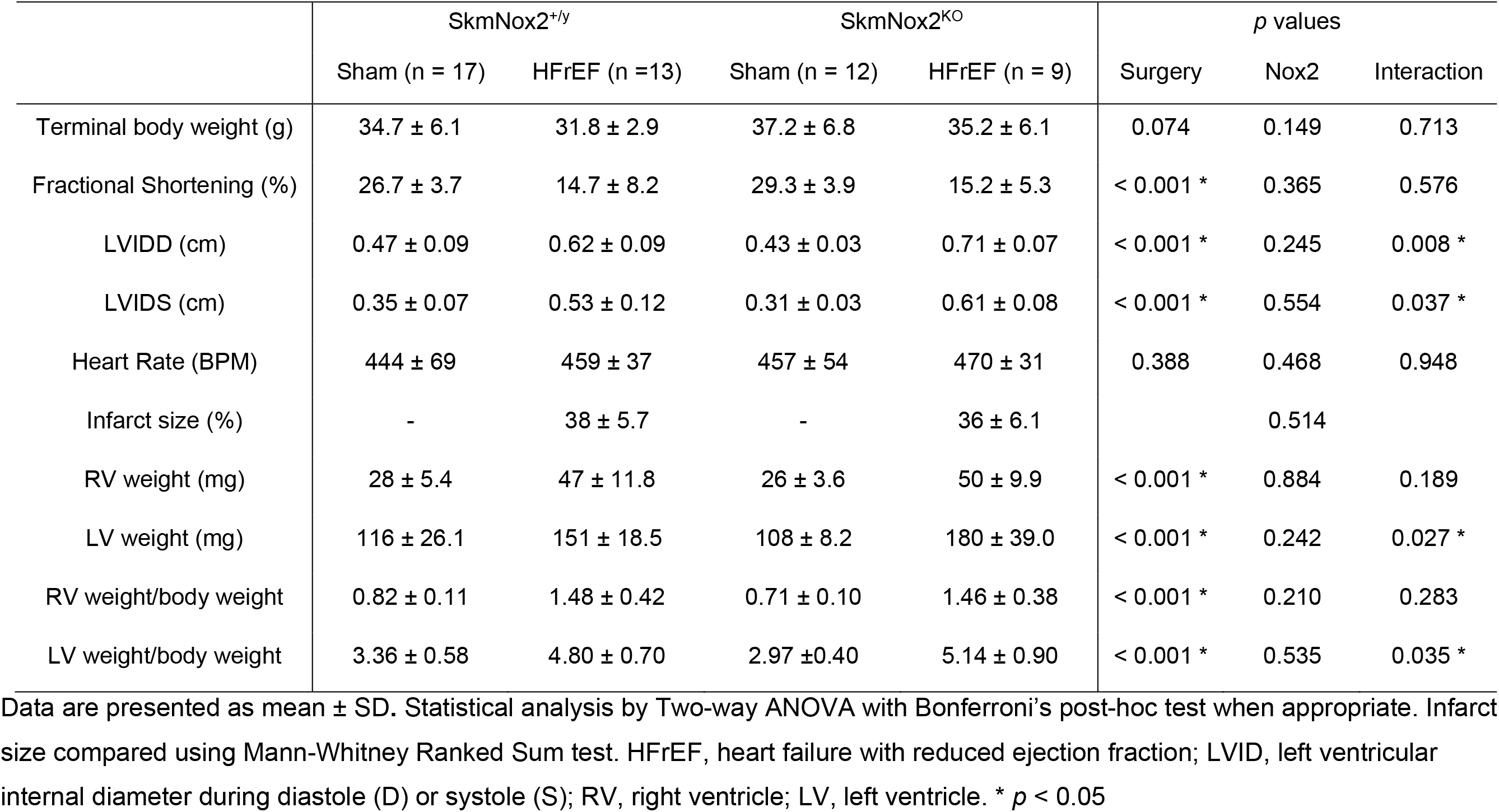
skmNox2 - Animal characteristics.

For investigations into the effect of skmNox2 on diaphragm function in HFrEF, a total of 48 animals underwent the MI procedure. skmNox2^KO^ mice had a 30% higher post-operative survival than skmNox2^+/y^ (*p* = 0.026) (Fig. 1B). After applying infarct size and right ventricle/body weight ratio inclusion criteria for severe HFrEF, thirteen skmNox2^+/y^ (one excluded) and nine skmNox2^KO^ (six excluded) HFrEF animals were included in final analyses. Seventeen skmNox2^+/y^ and twelve skmNox2^KO^ animals comprised the sham surgery groups. The total number of animals used for diaphragm analysis was n = 51. Ventricular weights and fractional shortening displayed typical signs of HFrEF in infarcted animals compared to controls in both skmNox4 and 2 cohorts (Tables 1 and 2).

### Diaphragm contractile properties

In the skmNox4 cohort, HFrEF caused a 20% drop in maximal specific force, but this was fully prevented by skmNox4 KO (Fig. 2A). There were significant surgery effects for HFrEF to decrease submaximal isometric contractions (twitch, Fig. 2B and 50 Hz, Table S1) with no protection by skmNox4 KO. There was a strain effect on 50 Hz contractions, with forces being higher in skmNox4KO (Table S1). Persistent reductions in submaximal force with HFrEF despite Nox4 KO suggested dysfunctional calcium handling which prompted assessment of the primary calcium release channel in skeletal muscle, the Ryanodine Receptor (RyR1). RyR1 content measured via immunoblotting decreased in both HFrEF groups with no effect from skmNox4 KO (Fig. 2G).

**Figure 2.**
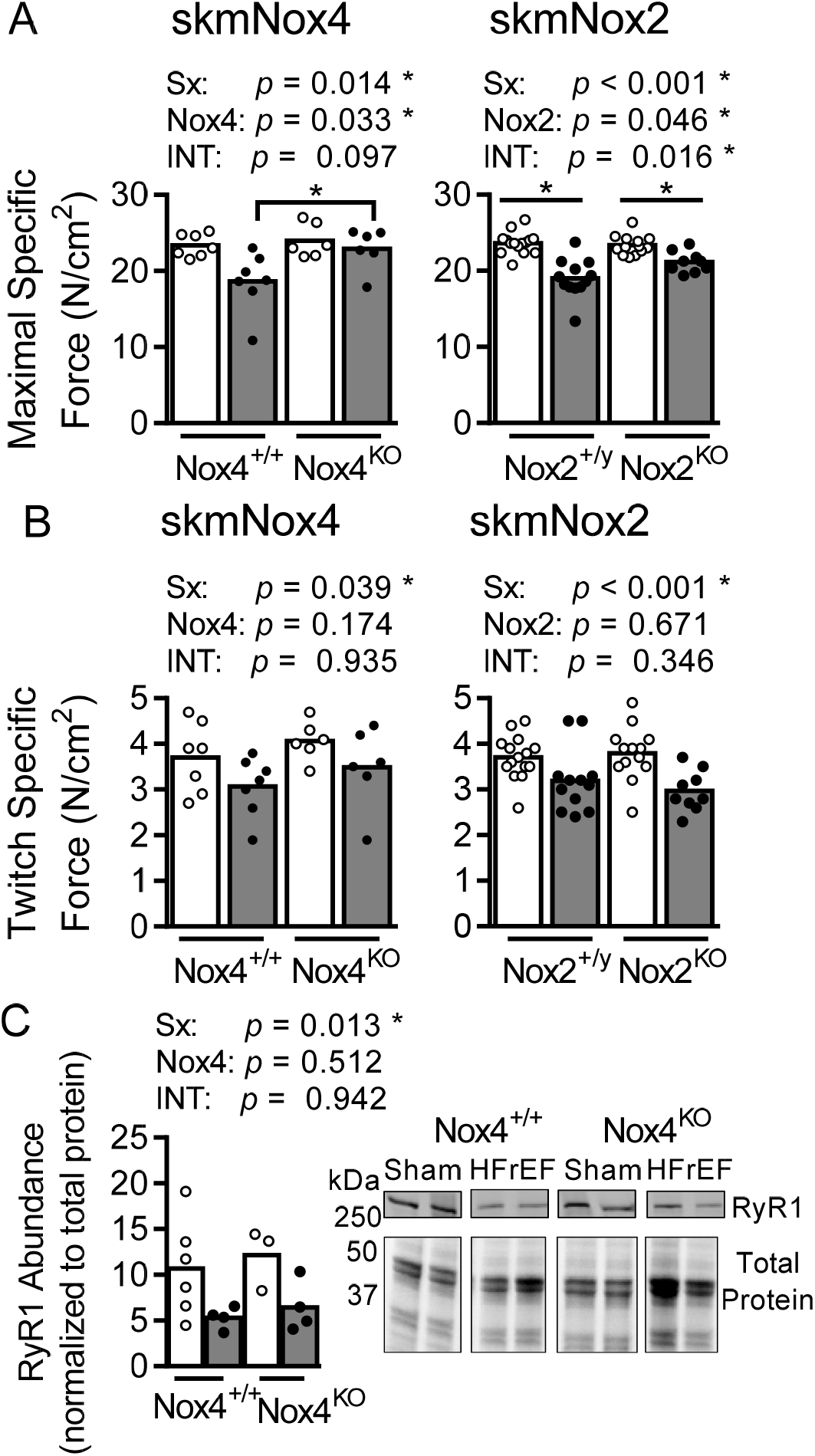
Diaphragm isometric contractile properties and RyR abundance. Maximal specific force (A) and twitch specific force (B) for skmNox4 and skmNox2 groups. (D) Diaphragm RyR abundance normalized to total protein (per lane). Images are representative membrane and total protein gel. Statistical analysis by two-way ANOVA with Bonferroni’s post-hoc test where appropriate. \**p* < 0.05

In the skmNox2 cohort, we found that HFrEF caused a 15% decline in maximal specific force in skmNox2^+/y^, while only a 7% drop for skmNox2^KO^ animals (Fig. 2A). Comparing maximal specific force between HFrEF groups showed a significant difference for skmNox2^+/y^ and skmNox2^KO^ (*p* = 0.041), which was not seen for Sham groups (*p* > 0.999). This shows that skmNox2^KO^ partially protected the diaphragm from contractile dysfunction. Similar to the Nox4 cohort, there were significant surgery effects for HFrEF to decrease submaximal isometric forces (twitch, 30 Hz, and 50 Hz) (Fig. 2B and Table S2) regardless of strain.

### Fiber type analysis and CSA

Animals in the Nox4 arm of experiments displayed no changes in cross sectional area or fiber type shifts with HFrEF or Nox4 KO (Fig. S3A, B). In the Nox2 arm of experiments, there were no observable fiber type shifts (Fig. S4D), but the combination of skmNox2^KO^ and HFrEF prompted a decline in the percentage of type IIa fibers and an increase in Type IIx fibers compared to skmNox2^KO^ shams (Fig. S4E).

### Mitochondrial respiration, content, and oxidant emission

Knockout of Nox4 increased mitochondrial *J*O_2_ under conditions of submaximal ADP and Complex I+II supported state 3 respiration (Fig.3A). There were no observable strain or surgery effects for Respiratory control ratio (RCR) (skmNox4-sham: 6.4 ± 1.6; skmNox4-HFrEF: 6.1 ± 1.4; skmNox4^KO^-sham: 6.6 ± 2.1; skmNox4^KO^-HFrEF: 6.2 ± 1.5), citrate synthase abundance, or mitochondrial ETC complexes subunits (Fig. 4B).

**Figure 3.**
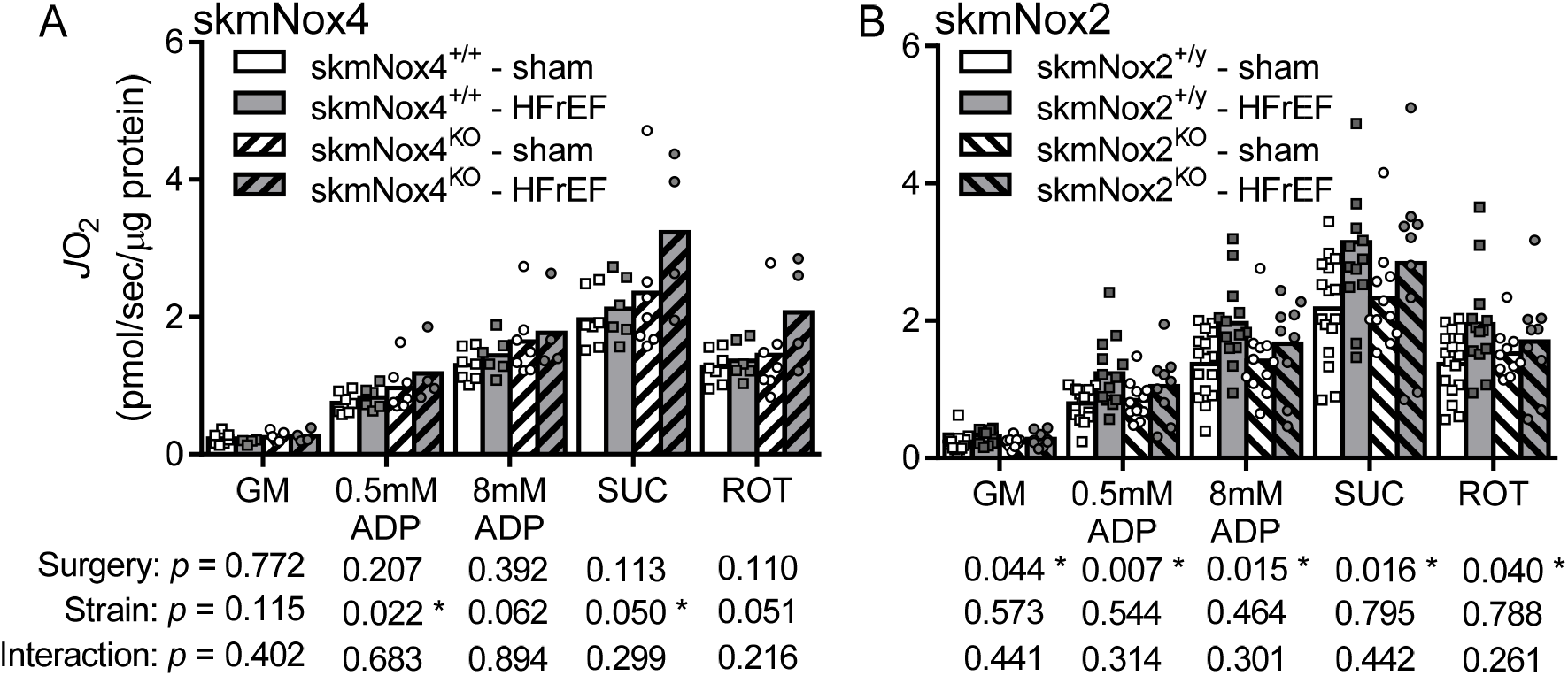
Mitochondrial Respiration. Rate of oxygen consumption (*J*O_2_) measured in saponin-permeabilized diaphragm bundles for (A) skmNox4 and (B) skmNox2 groups. Result normalized to fiber bundle protein content (*J*O_2_: pmol/sec/μg protein). Statistical analysis by two-way ANOVA for each substrate provided, with Bonferroni’s post-hoc test where appropriate. \**p* < 0.05

**Figure 4.**
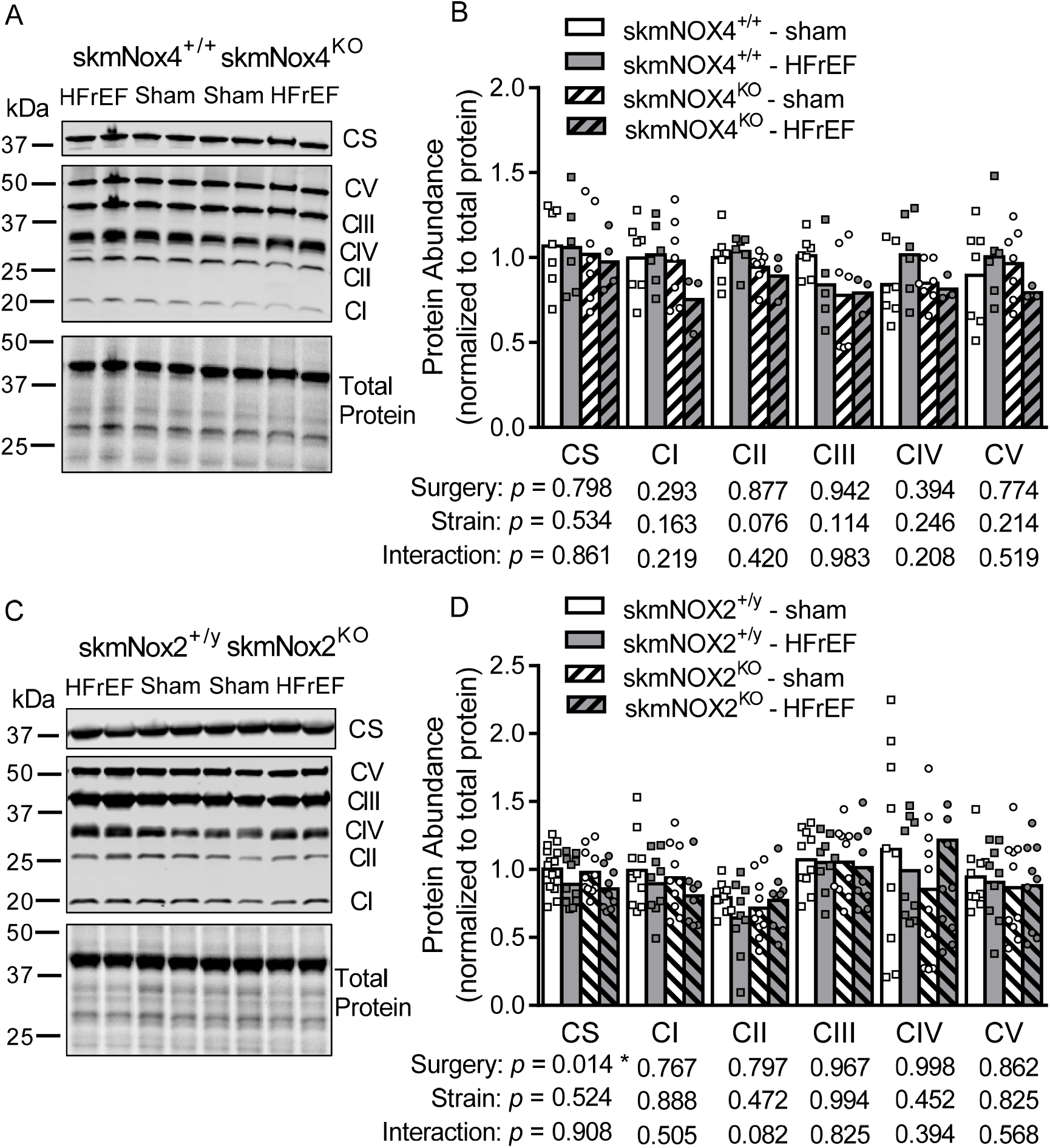
Diaphragm mitochondrial protein abundance. Representative membrane and gel images (A, C) and quantification for abundance of mitochondrial proteins (B, D) for skmNox4 (A, B) and skmNox2 (C, D) cohorts. Statistical analysis by two-way ANOVA. \**p* < 0.05. CS: citrate synthase; CI-V: mitochondrial complex I-V.

In the skmNox2 cohort, there was a significant surgery effect for HFrEF to increase mitochondrial *J*O_2_ under all conditions tested (Fig. 3B). RCR was not significantly different between groups (skmNox2-sham: 6.3 ± 2.0; skmNox2-HFrEF: 6.3 ± 1.4; skmNox2^KO^-sham: 6.4 ± 1.9; skmNox2^KO^-HFrEF: 6.1 ± 1.6). We also observed a significant surgery effect to decrease diaphragm citrate synthase content in HFrEF (Fig 4D), while there were no apparent changes in abundance of subunits of mitochondrial electron transport system complexes measured via immunoblotting (Fig. 4D).

Mitochondrial H_2_O_2_ emission showed no significant differences in either the Nox4 or Nox2 cohort for the conditions tested (Fig. 5A, B).

**Figure 5.**
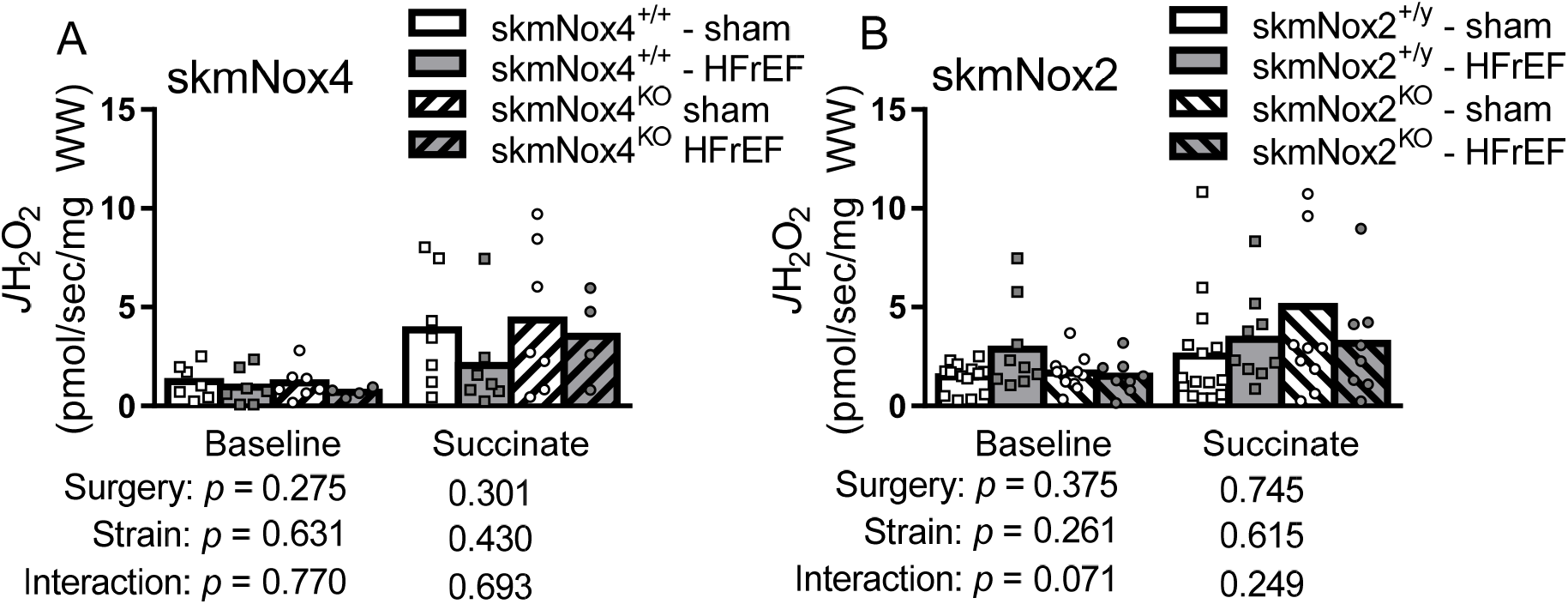
Mitochondrial H_2_O_2_ emission. Rate of hydrogen peroxide emission (*J*H_2_O_2_) for (A) skmNox4 and (B) skmNox2 groups. Statistical analysis by two-way ANOVA for condition. \**p* < 0.05.

### Glutathione content

We assessed total and oxidized glutathione content in the Nox4 cohort upon finding full protection of maximal specific force. Oxidized glutathione increased with HFrEF (skmNox4-sham: 1.10 ± 0.24; skmNox4-HFrEF: 1.47 ± 1.12; skmNox4^KO^-sham: 1.10 ± 0.47; skmNox4^KO^-HFrEF: 2.34 ± 1.43) (*p* = 0.039), while there were no statistically significance changes to GSH content (skmNox4-sham: 2.12 ± 0.85; skmNox4-HFrEF: 1.88 ± 0.75; skmNox4^KO^-sham: 1.96 ± 0.81; skmNox4^KO^-HFrEF: 3.27 ± 2.5)

## Discussion

The principal findings from this study are that skeletal muscle-specific knockout of NAD(P)H Oxidase 4 (Nox4) provided full protection against the loss of diaphragm maximal specific force in HFrEF, whereas skeletal muscle-specific knockout of Nox2 incurred partial protection of maximal specific force. Neither Nox4 nor Nox2 skeletal muscle-specific knockout protected against the loss of submaximal force. Our study did not reveal increased mitochondrial ROS or diminished respiration as potential mechanism for diaphragm weakness and protection by Nox4 or Nox2. In fact, mitochondrial respiration was increased in diaphragm bundles from animals with HFrEF in the skmNox2 arm of the study. Knockout of Nox4 increased mitochondrial respiration independent of surgery (sham vs. MI). Finally, knocking out Nox2 from skeletal myocytes increased survival post-myocardial infarction. These findings highlight a critical role for skeletal myocyte NAD(P)H Oxidases to influence maximal specific force (Nox4 and Nox2) and systemic pathophysiology (Nox2) in response following myocardial infarction.

### Mouse model of HFrEF

Myocardial infarction (MI) is an established model of HFrEF with translational relevance (Bayat *et al*., 2002; Patten & Hall-Porter, 2009). The MI procedure has a survival rate of ~50%, with most deaths occurring within 3-10 days (Bayat *et al*., 2002; Gao *et al*., 2005; Gao *et al*., 2010). This agrees with guidelines for experimental models of myocardial ischemia and infarction (Lindsey *et al*., 2018) and in line with what we observed for the skmNox2^+/y^, skmNox4^+/+^ and skmNox4^KO^ MI groups. Interestingly, knockout of Nox2 in skeletal muscle cells increased survival post-MI to ~80% while infarct size was similar across MI groups. This suggests a role for skeletal myocyte Nox2 to influence the systemic pathophysiology post-myocardial infarction, which could be possible through inflammatory signaling via skeletal myokines that modulate cardiac remodeling and the development of HFrEF. For instance, Nox2 modulates the release of IL-6 from skeletal myotubes (Henríquez-Olguín *et al*., 2015). Our study was not designed to examine survival post-MI and its related mechanisms - the focus was on the chronic stage of HFrEF, and increased survival was a serendipitous finding that warrants future investigation.

Because of the relationship between diaphragm weakness and disease severity (Kelley *et al*., 2020), we limited our analyses to animals exhibiting signs of severe HFrEF based on RV weight-to-body weight ratio and infarct size. These criteria resulted in a greater number of animals excluded from the skmNox2^KO^ group (n = 6) than the skmNox2^+/y^ group (n = 1), further supporting the hypothesis that skeletal muscle Nox2 can influence cardiac remodeling postmyocardial infarction. In the Nox4 cohort, a similar number of animals was excluded from the MI groups. Per inclusion criteria, animals included in final analyses displayed typical signs of severe HFrEF (left and right ventricular hypertrophy, reduced fractional shortening, and a large infarcted area) that were not different following genetic recombination (KO vs. +/+).

### Diaphragm weakness – contractile function

Diaphragm weakness is common in patients with severe HFrEF (Mancini *et al*., 1995; Kelley & Ferreira, 2017; Miyagi *et al*., 2018). The diaphragm is involved in inspiration and airway clearance maneuvers, and weakness in this muscle has been associated with the key HFrEF symptoms of dyspnea, fatigue, and exercise intolerance (Kelley & Ferreira, 2017).

Diaphragm weakness can occur independent of muscle atrophy and is believed to be caused by excess production of reactive oxygen species from unknown sources (Kelley & Ferreira, 2017). The decreases in diaphragm specific force observed in the HFrEF groups of this study (~20% for maximal specific force and twitch contractions) align with previous reports of diaphragm weakness in HFrEF (Ahn *et al*., 2015; Laitano *et al*., 2016; Adams *et al*., 2019; Kelley *et al*., 2020). Skeletal muscle specific knockout of the Nox2 subunit provided partial protection to maximal specific force, while the knockout of Nox4 incurred full protection. The lack of full protection in the skmNox2^KO^ animals suggests that Nox2 is not the only cause of diaphragm weakness, or that knockout of Nox2 results in compensatory responses from other sources of ROS. Indeed, a recent report found that knockout of the Nox2 assembly unit p47^phox^ leads to a compensatory increase in Nox4 (Cully & Rodney, 2020). Nox4 may then contribute to increased oxidant production with the knockout of Nox2. In skmNox2KO animals, mRNA abundance of Nox4 increased with HFrEF, but did not reach significance (*p* = 0.099). Regardless, Nox4 plays a pivotal role in the development of diaphragm weakness, as seen in our findings of full protection of maximal specific force with the knockout of this protein.

The observation of consistently impaired submaximal force (1, 30, 50 Hz) in HFrEF, even with skmNox2 or skmNox4 KO, points to impaired Ca^2+^ handling or Ca^2+^ sensitivity of the myofilaments. Based on these findings and evidence of RyR dysfunction in human diaphragm biopsies from Mangner et al. (2021), we followed up with immunoblots of the RyR1 and uncovered decreased RyR1 content in the diaphragm of HFrEF animals in the Nox4 cohort. The RyR1 is the primary calcium release channel in skeletal muscle, releasing calcium from the sarcoplasmic reticulum into the cytosol to initiate muscle contraction. Other experiments in the diaphragm have found evidence of impaired Ca^2+^ handling with HFrEF (Dominguez & Howell, 2003; Mangner *et al*., 2021) as well as reports of a rightward shift in the force-pCa^2+^ curve of isolated diaphragm muscle fibers, indicating reduced calcium sensitivity (van Hees *et al*., 2007; Hwee *et al*., 2015; Mangner *et al*., 2021). In limb skeletal muscle, hyperphosphorylation of RyR1 impairs calcium handling and muscle contractile function in HFrEF (Wehrens *et al*., 2005). RyR1 dysfunction and decreased abundance seems to occur independent of Nox2 and Nox4. It is also unlikely that Nox2 or Nox4 exerted any protection against myofilament desensitization to calcium suggesting that mechanisms unrelated to ROS are responsible for that process.

### Diaphragm muscle fiber type and cross-sectional area

Another potential contributor to diaphragm weakness in HFrEF is muscle atrophy (Howell *et al*., 1995; Adams *et al*., 2019; Kelley *et al*., 2020; Mangner *et al*., 2021), but this is not a consistent finding in rodents (van Hees *et al*., 2007; Laitano *et al*., 2016). Diaphragm atrophy is associated with activation of proteolytic pathways, and inhibiting the ubiquitin-proteasome pathway was sufficient to protect against both diaphragm atrophy and loss of force in response to HFrEF (van Hees *et al*., 2008; Adams *et al*., 2019). We did not observe diaphragm atrophy in the HFrEF groups of either the Nox2 or Nox4 arms of the study. This discrepancy compared to other reports is possibly due to our mouse model of HFrEF, as multiple reports in larger mammals have observed diaphragm muscle atrophy in response to HFrEF (Howell *et al*., 1995; Dominguez & Howell, 2003; Kelley *et al*., 2020). Other explanations may be related to mouse sex (atrophy observed in female mice (Adams *et al*., 2019)) or strain (atrophy observed in CD-1 mice (Gillis *et al*., 2016). While atrophy has been reported in previous studies and occurs in humans, it does not contribute to the diaphragm weakness we observed herein.

Fiber type shifts have also been observed in diaphragm biopsies of patients with HFrEF; unlike the increase in glycolytic Type II fibers that is observed in limb muscle (Adams *et al*., 2017), the diaphragm experiences a shift to increased type I fibers and diminished Type II fibers (Tikunov *et al*., 1996; Tikunov *et al*., 1997; Mangner *et al*., 2021). This is believed to result from the increased work and frequency of breathing that occurs in patients with severe HFrEF, leading to an oxidative phenotype in the muscle fibers that is akin to endurance exercise training. Coupled with these fiber type shifts, there are several reports of increased mitochondrial-associated enzymes in humans and animal models of HFrEF (Howell *et al*., 1995; Tikunov *et al*., 1997; Laitano *et al*., 2016; Mangner *et al*., 2021). Shifts in fiber type as seen in patients with HFrEF may not occur or may be more difficult to observe in the mouse diaphragm because of its homogenous distribution of muscle fibers (90% type II, 10% type I (Adams *et al*., 2019)), despite its relatively high mitochondrial content (Hahn *et al*., 2019). Importantly, shifts in myosin heavy chain isoform, which are commonly used to delineate fiber type, do not always reflect the shifts in mitochondrial content and oxidative capacity.

### Mitochondrial function

For assessments of mitochondrial function, we measured mitochondrial respiration in saponin-permeabilized diaphragm bundles. In the Nox2 cohort, we found an increase in maximal mitochondrial respiration (state 3 with complex I and II substrates) with HFrEF independent of Nox2, but no differences in respiratory control ratio (state 3/state 2 respiration). These coupled findings suggest an increase in diaphragm mitochondrial density without a change in mitochondrial respiratory function. However, when we probed for changes in mitochondrial content via immunoblotting, we observed a contradictory decrease in citrate synthase abundance in the HFrEF groups and no change in mitochondrial complexes.

In the skmNox4 cohort, ADP-stimulated respiration was significantly higher with the knockout of Nox4, indicating a critical role for this protein on mitochondrial function. Nox4 is localized to the mitochondria (Ago *et al*., 2010; Beretta *et al*., 2020), and knockdown of Nox4 in other cell types increases respiration and markers of mitochondrial content (Kozieł *et al*., 2013; Bernard *et al*., 2017). Surgery effects on respiration may have been masked by effects of Nox4 KO on diaphragm mitochondria or due to a smaller sample size. Similar to findings in Nox2 KO and controls, there were no changes in respiratory control ratio which suggested a change in mitochondrial content with Nox4 knockout. This was not supported by immunoblotting of citrate synthase and mitochondrial complexes, which showed no changes in protein abundance based on either surgery or genetic strain.

The gold standard for assessing mitochondrial function and content in skeletal muscle are respirometry and mitochondrial fractional area measured via electron microscopy (Larsen *et al*., 2012). Several surrogate metrics (enzyme content and activity assays) correlate well with electron microscopy in healthy young men (Larsen *et al*., 2012), but there are discrepancies that may be further exacerbated in diseases such as HFrEF. Critically, no protein appears to be a uniform marker of mitochondrial content across different organs (McLaughlin *et al*., 2020) but citrate synthase activity provides the best approximation within skeletal muscle (Larsen *et al*., 2012). All considered, our respirometry findings of increased oxidative capacity in the diaphragm likely offer the best explanation of mitochondrial adaptations that occur in response to HFrEF (skmNox2 cohort) or Nox4 KO (skmNox4 cohort) in the current study. Studies in humans have noted increased citrate synthase activity in diaphragm biopsies from patients with HFrEF (Tikunov *et al*., 1997; Mangner *et al*., 2021), indicating increased mitochondrial content that matches the increase in Type I muscle fibers. However, this increase in content is coupled with decreased mitochondrial respiration and a decrease in mitochondrial size in electron microscopy (Mangner *et al*., 2021). Importantly, changes in fiber type and increased mitochondrial function do not result in functional improvements in the muscle, as inspiratory muscle fatigue is reported in patients with HFrEF (Mancini *et al*., 1995).

### Reactive oxygen species and redox balance

Diaphragm weakness in HFrEF is linked to excessive production of ROS that oxidize myofibrillar and calcium handling proteins in skeletal muscle. In the skmNox4 cohort, we observed a significant increase in oxidized glutathione measured in diaphragm muscle homogenates of HFrEF animals, accompanied by no change in reduced glutathione. This aligns with several previous reports of increased markers of oxidized shifts in redox balance in the diaphragm of HFrEF patients and animal models (Supinski & Callahan, 2005; Ahn *et al*., 2016; Kelley *et al*., 2020; Mangner *et al*., 2021), but the exact source of increased ROS or oxidized shift has not been resolved. Diaphragm Nox2 and Nox4 abundance increases in response to HFrEF (Ahn *et al*., 2015; Ahn *et al*., 2016; Mangner *et al*., 2021), and there is increased phosphorylation of the Nox2 assembly subunit p47phox (Ahn *et al*., 2015), implicating Nox2 and Nox4 as possible sources of excess ROS.

Nox4 expression increased with HFrEF, and this was prevented with our targeted Nox4 KO. However, we did not observe increased expression of Nox2 with HFrEF. Expression of Nox proteins may go through compensatory and temporal changes over the 16-week post-MI and in response to manipulation redox-related enzymes. A recent report that whole-body knockout of p47phox) presumably preventing ROS from Nox2) lead to a compensatory increase in abundance of Nox4 in skeletal muscle (Cully & Rodney, 2020). This certainly appears to be the case in our study as well, as knockout of Nox4 lowered mRNA abundance of Nox2 and p47phox and knockout of Nox2 led to an increase in Nox4 (though not reaching the threshold for statistical significance). Importantly, our measurements of Nox2 and 4 mRNA were made with whole-muscle homogenates that contain other cell types, including vascular, neural, and immune cells that may influence our results.

A previous study from our lab reported that whole-body knockout of the Nox2 complex organizing subunit p47^phox^ provides full protection of diaphragm maximal and submaximal force in the same mouse model of HFrEF employed here (Ahn *et al*., 2015). This contrasts with the findings of partial protection of maximal force and no protection of submaximal force with the skeletal muscle-specific knockout of the Nox2 subunit. There are several possibilities for this discrepancy: 1) whole-body knockout of p47^phox^ prevents the development of severe HFrEF and consequent diaphragm weakness. This appears unlikely because p47^phox^ KO animals from the aforementioned study show signs of more severe HFrEF compared to controls; 2) p47^phox^ interacts with and influences Nox4 activity. While there are no reports of Nox4 associating with cytosolic subunits such as p47^phox^, Nox4 does contain a p47^phox^ binding motif (von Löhneysen *et al*., 2010). p47^phox^ KO could then prevent ROS production from both Nox2 and Nox4 leading to full protection of maximal and submaximal force in HFrEF; or 3) other cell types (i.e., vascular, neuronal, and immune) with high Nox2 activity contribute to diaphragm weakness, and these are still active with a skeletal muscle-specific intervention. This seems the most probable considering the full protection of maximal specific force with Nox4 KO.

The mitochondrion is another skeletal muscle source of ROS, which may increase following cross-talk with other ROS sources (Dikalov, 2011). Previous studies in rats have observed increase mitochondrial H_2_O_2_ emission following myocardial infarction (Supinski & Callahan, 2005; Laitano *et al*., 2016; Coblentz *et al*., 2019). Our methods to measure mitochondrial H_2_O_2_ with permeabilized mouse diaphragm fibers followed a similar approach as the aforementioned studies but there were no significant differences due to HFrEF. This suggests either that mouse diaphragm mitochondria undergo differing adaptations to the rat, or that the assay in mouse diaphragm fibers may not be sensitive enough to detect differences between sham and HFrEF.

Finally, xanthine oxidase (XO) is an additional source of ROS within skeletal muscle that is worth mentioning in the context of muscle weakness in HFrEF. XO content increases in the early stage (72 hr.) of HFrEF in both diaphragm and limb muscle (Scott Bowen *et al*., 2015; Nambu *et al*., 2021). However, XO activity is not elevated long term in limb muscle (days 7-28 post MI) and diaphragm XO content has not been explored in chronic HFrEF. One interpretation is that the mechanisms of muscle weakness are different in the early (up to 72 hours) vs. late/chronic stage of HFrEF, with XO playing predominant role in the early phase. Indeed, a recent study from our lab investigated diaphragm weakness 72 hours post-MI and found no protection with whole-body KO of Nox4 (Hahn *et al*., 2021), contrasting with our findings with skmNox4 KO in chronic HFrEF.

### Limitations

There are several limitations to the current work. First, we did not measure Nox2 or 4 activities in skeletal muscle. Chemiluminescence assays have been used as a proxy for Nox activity, but their specificity has been questioned because experiments using a Nox1 −2 and −4 triple knockout did not observe changes in signal (Rezende *et al*., 2016). A reliable Nox activity assay that is validated by appropriate positive and negative controls has not yet been developed Finally, our method to measure mitochondrial H_2_O_2_ emission and analyze crosstalk between Nox and mitochondrial-derived ROS was limited to reverse electron transport and ROS emission from Complex I following the addition of succinate. Nox4 localizes in the mitochondria and is capable of interacting with Complex I (Hirschhäuser *et al*., 2015), suggesting Complex I as a critical source of mitochondrial-derived ROS in HFrEF and Nox-crosstalk. While we did not observe changes in succinate-induced ROS emission in HFrEF or with skmNox4 or 2 KO, there are several additional sites of ROS production within the mitochondria that cannot be excluded (Murphy, 2009).

## Conclusions

Findings from our study indicate that Nox4 and Nox2 serve critical roles in the loss of diaphragm maximal specific force with HFrEF. Maximal inspiratory pressure, a measure of maximal diaphragm strength in humans, predicts adverse outcomes in patients with HFrEF. Thus, targeting skeletal muscle Nox4 or 2 could improve diaphragm strength and outcomes in patients with HFrEF. In our study, diaphragm weakness was not related to changes in muscle fiber type or atrophy, and succinate induced mitochondrial H_2_O_2_ emission did not change in response to our genetic or surgical interventions. Mitochondrial respiration increased with HFrEF but occurred in tandem with decreases or no change in markers of mitochondrial content, suggesting a mismatch between mitochondrial content and function that has also been observed in humans. Finally, the knockout of Nox2 specifically from skeletal muscle increased survival post-myocardial infarction indicating a crucial role for skeletal muscle on the systemic pathophysiology following myocardial infarction.

## Acknowledgments

The study was funded by NIH R01 HL1331806. R. Kumar received a predoctoral fellowship from the American Heart Association (20PRE35200047). L. Ferreira was supported by a University of Florida Research Foundation Professorship. T. Ryan was funded by NIH R01 HL149704.

