## Supplemental figures for "Skeletal muscle Nox4 knockout prevents and Nox2 knockout blunts loss of maximal diaphragm force in mice with heart failure with reduced ejection fraction"

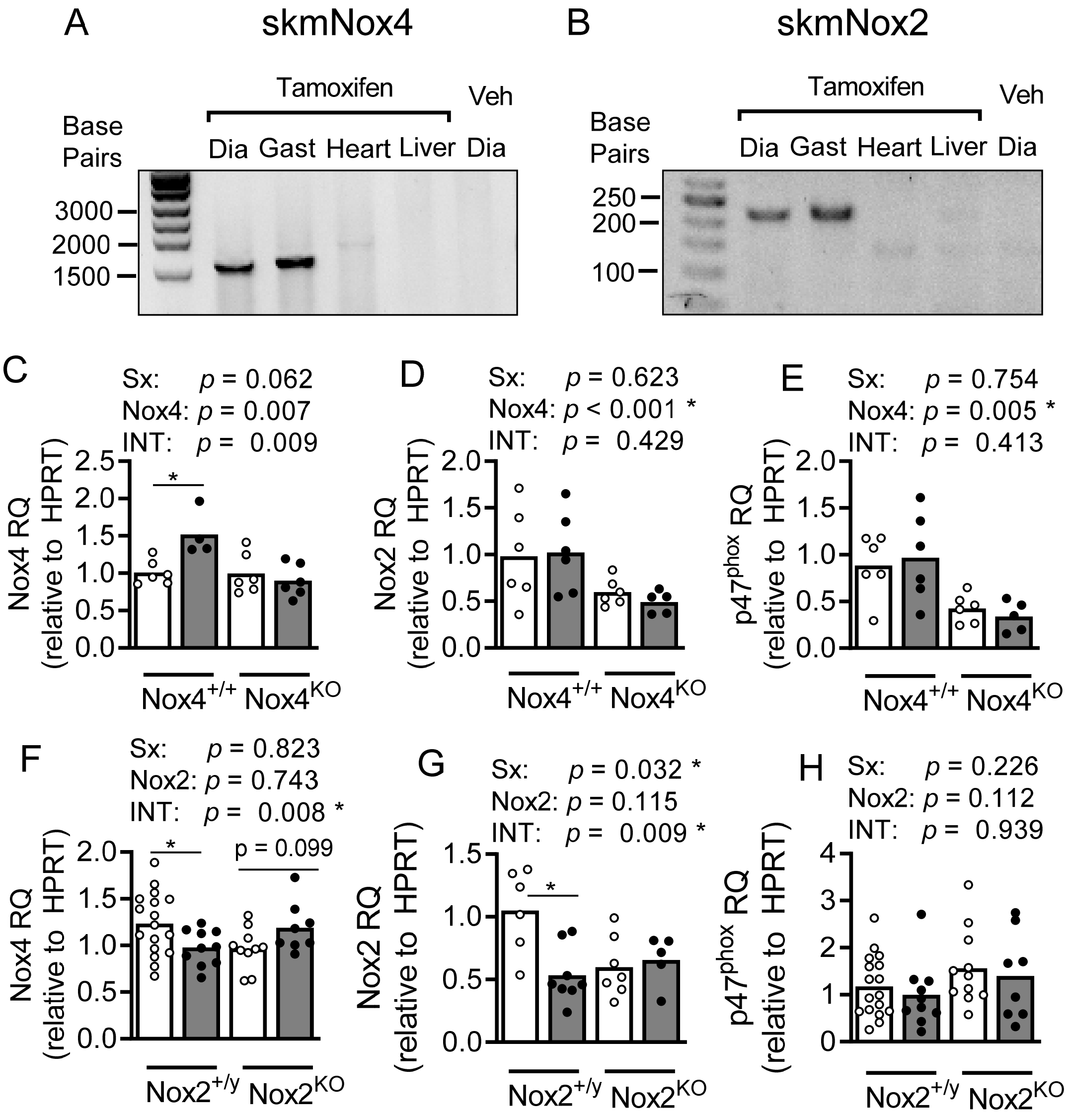


**Supplemental Fig. 1 – Confirmation of Cre-induced genetic recombination and mRNA abundance of Nox subunits.** (A, B) Representative gel images confirming genetic recombination in the diaphragm and gastrocnemius muscles of experimental animals receiving tamoxifen injections. Amplification of ~1600 base pair (bp) product in skmNox4 animals (A) and 225 bp product in skmNox2 animals (B). No recombination observed in the heart or liver of tamoxifen injected animals, or in the diaphragm from animals receiving control/vehicle injections. mRNA abundance of Nox4 (C,F), Nox2 (D, G), and $p47^{phox}$ (E, H) measured from diaphragm muscle homogenates for the skmNox4 (C, D, E) and skmNox2 (F, G, H). Statistical analysis by two-way ANOVA with Bonferroni's post-hoc test where appropriate. $^{*}p < 0.05$.

**Supplemental Table 1 – skmNox4 diaphragm contractile properties**

| | SkmNox4$^{+/+}$ | | SkmNox4KO | | *p* values | | |
|---|---|---|---|---|---|---|---|
| | Sham (n = 7) | HFrEF (n = 7) | Sham (n = 7) | HFrEF (n = 7) | Surgery | Nox4 | Interaction |
| 30 Hz Specific force (N/cm$^2$) | 5.61 ± 1.70 | 4.71 ± 1.27 | 6.18 ± 0.69 | 5.45 ± 1.93 | 0.174 | 0.274 | 0.887 |
| 50 Hz Specific force (N/cm$^2$) | 12.10 ± 2.47 | 9.76 ± 1.21 | 13.88 ± 1.70 | 12.32 ± 3.51 | 0.046 * | 0.028 * | 0.678 |
| Max Rate of Contraction (% Max Force/ms) | 3.24 ± 0.40 | 3.14 ± 0.53 | 3.17 ± 0.33 | 2.99 ± 0.34 | 0.401 | 0.524 | 0.820 |
| Max Rate of Relaxation (% Max Force/ms) | -5.23 ± 0.55 | -5.09 ± 0.58 | -5.12 ± 0.57 | -4.88 ± 0.50 | 0.409 | 0.495 | 0.829 |
| Time to peak tension (ms) | 15.14 ± 1.21 | 14.50 ± 1.05 | 15.83 ± 0.75 | 14.83 ± 1.60 | 0.101 | 0.298 | 0.713 |
| ½ Relaxation Time (ms) | 14.26 ± 2.24 | 16.13 ± 3.30 | 14.37 ± 1.46 | 15.92 ± 1.88 | 0.080 | 0.957 | 0.865 |

Data are presented as mean ± SD**.** Statistical analysis by Two-way ANOVA with Bonferroni's post-hoc test when appropriate. * $p < 0.05$

**Supplemental Table 2 – skmNox2 diaphragm contractile properties**

| | SkmNox2$^{+/y}$ | | SkmNox2$^{KO}$ | | *p* values | | |
|---|---|---|---|---|---|---|---|
| | Sham (n = 7) | HFrEF (n = 7) | Sham (n = 7) | HFrEF (n = 7) | Surgery | Nox2 | Interaction |
| 30 Hz Specific force (N/cm$^2$) | 5.73 ± 0.84 | 5.15 ± 1.29 | 5.82 ± 0.82 | 4.48 ± 0.76 | 0.001 * | 0.294 | 0.181 |
| 50 Hz Specific force (N/cm$^2$) | 12.88 ± 1.95 | 10.43 ± 1.88 | 13.42 ± 1.20 | 10.30 ± 1.97 | < 0.001 * | 0.686 | 0.518 |
| Max Rate of Contraction (% Max Force/ms) | 3.07 ± 0.28 | 3.28 ± 0.54 | 3.09 ± 0.57 | 2.99 ± 0.29 | 0.939 | 0.563 | 0.174 |
| Max Rate of Relaxation (% Max Force/ms) | -5.16 ± 0.35 | -4.97 ± 0.55 | -5.18 ± 0.36 | -5.14 ± 0.43 | 0.200 | 0.248 | 0.307 |
| Time to peak tension (ms) | 15.13 ± 1.25 | 15.42 ± 2.57 | 15.92 ± 1.19 | 15.00 ± 1.87 | 0.535 | 0.717 | 0.245 |
| ½ Relaxation Time (ms) | 14.12 ± 1.83 | 15.25 ± 2.45 | 13.43 ± 1.34 | 14.35 ± 1.03 | 0.054 | 0.131 | 0.839 |

Data are presented as mean ± SD**.** Statistical analysis by Two-way ANOVA with Bonferroni's post-hoc test when appropriate. * $p < 0.05$

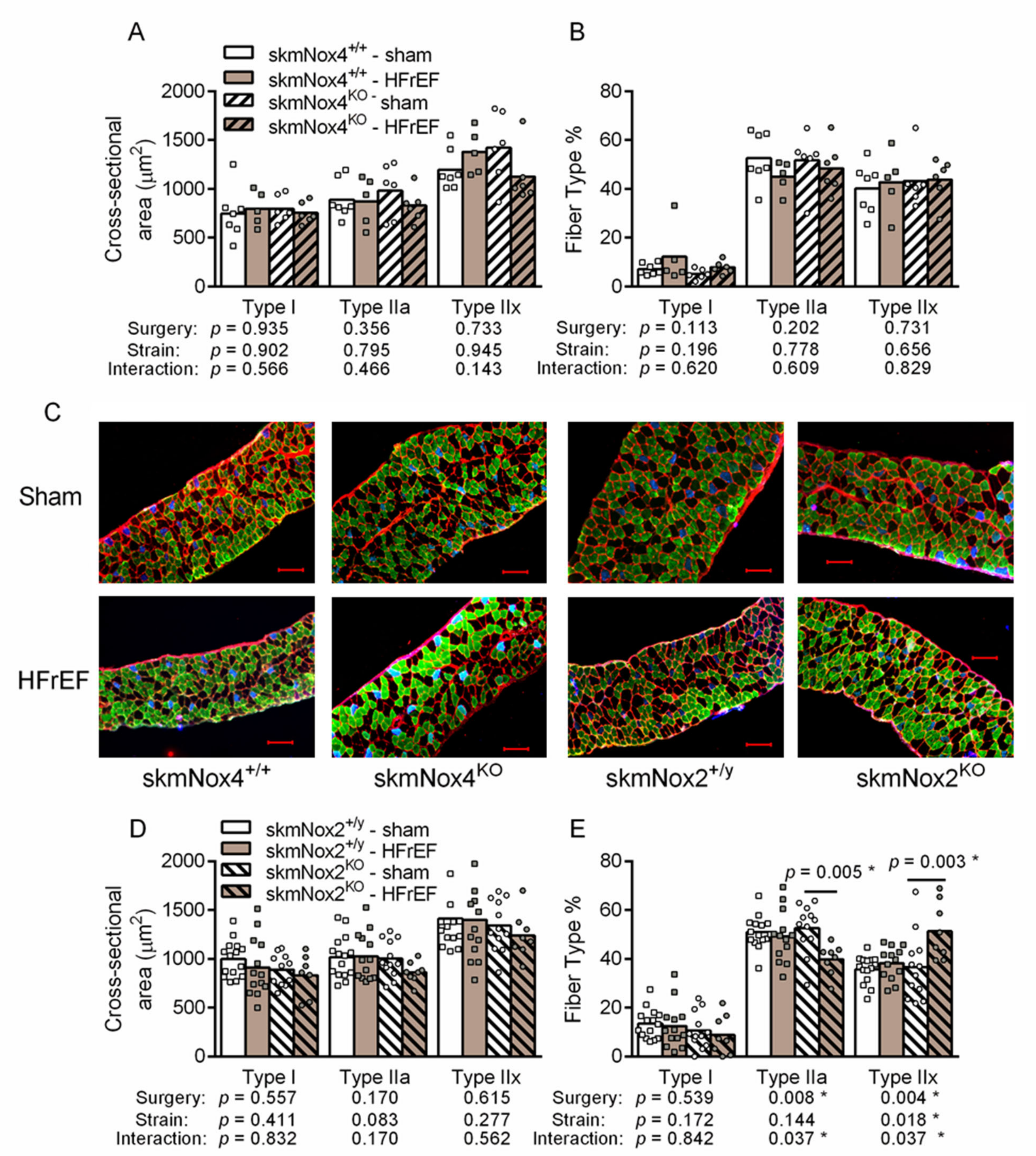


**Supplemental Fig. 3 – Diaphragm cross-sectional area and muscle fiber type distribution.** Fiber cross-sectional area by fiber type (A, C) and fiber type frequency (%) (B, D) from diaphragm micro sections. Scale bars represents 100 µM. Colors represent specific myosin heavy chain (MyHC) isoforms (blue = type I, green = type IIa, black = type IIb/x). Statistical analysis of cross-sectional area by linear mixed modeling. Statistical analysis of fiber type by two-way ANOVA with Bonferroni's post-hoc test where appropriate. *$p < 0.05$
